# NAD^+^ treatment can increase directly the antioxidant capacity of rotenone-treated differentiated PC12 cells

**DOI:** 10.1101/498162

**Authors:** Jie Zhang, Yunyi Hong, Wei Cao, Haibo Shi, Weihai Ying

## Abstract

NAD^+^ administration can produce profound beneficial effects in the animal models of aging and a number of diseases. Since oxidative stress plays key pathological roles in aging and multiple major disorders, it is crucial to elucidate the mechanisms underlying the protective effects of NAD^+^ administration on oxidative stress-induced cell death. Previous studies have suggested that NAD^+^ treatment can decrease oxidative cell death indirectly by such mechanisms as preventing mitochondrial permeability transition, while it is unclear if NAD^+^ administration may decrease oxidative cell death by increasing directly the antioxidant capacity of the cells. Our current study used rotenone-treated differentiated PC12 cells as a cellular model to test our hypothesis that NAD^+^ treatment may increase directly the antioxidant capacity of the cells exposed to oxidative stress. Our study has indicated that NAD^+^ treatment can significantly attenuate the rotenone-induced increase in oxidative stress in the cells. Moreover, NAD^+^ treatment can significantly enhance the GSH/GSSG ratio - a major index of antioxidant capacity - of rotenone-treated cells. Collectively, our study has provided the first evidence indicating that NAD^+^ treatment can increase directly the antioxidant capacity of cells exposed to oxidative stress. These findings have suggested a novel mechanism underlying the profound protective effects of NAD^+^ administration in numerous disease models: NAD^+^ administration can decrease oxidative stress-induced cell death by enhancing directly the antioxidant capacity of the cells. Our finding has also highlighted the nutritional potential of NAD^+^.

## Introduction

NAD^+^ plays important roles in multiple biological functions, including energy metabolism, mitochondrial functions and immune functions [12, 15, 16]. NAD^+^ administration can profoundly decrease the tissue injury in animal models of numerous diseases and aging [18-20], suggesting that NAD^+^ deficiency may be a key common pathological factor of multiple major diseases and aging [19]. It is crucial to further investigate the mechanisms underlying the protective effects of NAD^+^ administration, which is required for developing NAD^+^ administration into an effective therapeutic strategy for multiple diseases.

Due to the critical pathological roles of oxidative stress in aging and numerous diseases [7, 11], it is of significance to investigate the mechanisms underlying the protective effects of NAD^+^ treatment on oxidative cell death. It has been reported that NAD^+^ treatment can decrease oxidative stress-induced death of neurons, astrocytes and myocytes [1, 2, 17, 20]. Several studies have suggested that NAD^+^ treatment decreases oxidative cell death indirectly by such mechanisms as prevention of glycolytic inhibition, mitochondrial permeability transition, mitochondrial depolarization and nuclear translocation of apoptosis-inducing factor (AIF) [1-3, 8].

In our current study, we used rotenone-treated differentiated PC12 cells as a cellular model to test our hypothesis that NAD^+^ treatment can decrease oxidative cell death by enhancing directly the antioxidant capacity of the cells. Our study has provided evidence supporting our hypothesis: NAD^+^ administration can significantly decrease rotenone-induced oxidative stress, as well as enhance the ratio of GSH/GSSG of the cells.

## Materials and Methods

### Cell cultures

Cell cultures were prepared as described previously [5]. Differentiated PC12 cells were purchased from the Cell Resource Center of Shanghai Institute of Biological Sciences, Chinese Academy of Sciences. The cells were plated onto 6-well or 24-well cell culture plates at the initial density of 1×10^5^ cells/ml in Dulbecco’s Modified Eagle Medium (DMEM) containing 4500 mg/L D-glucose, 584 mg/L L-glutamine, 110 mg/L sodium pyruvate (Thermo Scientific, Waltham, MA, USA), which also contains 1% penicillin and streptomycin (Invitrogen, Carlsbad, CA, USA) and 10% fetal bovine serum (PAA, Linz, Austria). The cells were used when the density of the cell cultures reached 60-80%.

### Experimental procedures

Experiments were initiated by replacing the cell culture medium with medium containing various concentrations of drugs. The cells were left in an incubator with 5% CO_2_ at 37 °C for various durations.

### FACS-based ethidium fluorescence assay

FACS-based assays using dihydroethidium (DHE) as an oxidative stress probe were conducted to detect the superoxide levels in PC12 cells. DHE can be oxidized by superoxide to become ethidium that becomes fluorescent after it binds to DNA or RNA [4]. In brief, cells were washed with PBS, harvested, collected and incubated in 5 μM DHE probes diluted in serum-free DMEM at 37°C for 20 min. Then the cells were centrifuged, washed and re-suspended in PBS. The ethidium fluorescence was examined by FACS at excitation wavelength of 535 nm and emission wavelength of 590 nm. Ten thousands cells were counted for each sample.

### Determinations of GSH, GSSG, and GSH/GSSG ratios

The levels of total glutathione (GSH + GSSG) and GSSG were determined by using a commercially available kit (Beyotime Institute of Biotechnology, Haimen, China), which was conducted according to the manufacturer’s instructions. In brief, cells were washed with PBS for three times and lysed, then frozen at −80 °C and thawed at room temperature for three times. The levels of total glutathione were assessed by the 5, 5′ dithiobis (2-nitrobenzoic acid) (DTNB)-oxidized glutathione (GSSG) reductase-recycling assay. In brief, 700 μl of 0.33 mg/ml NADPH in 0.2 M sodium phosphate buffer containing 10 mM ethylenediaminetetraacetic acid (pH 7.2), 100 μl of 6 mM DTNB, and 190 μl of distilled water were added and mixed in an eppendoff tube. The reaction was initiated by addition of 10 μl of 250 IU/ml glutathione reductase. With readings recorded every 30 seconds, the absorbance was monitored at 412 nm by a plate reader for 15 minutes. To determine the levels of GSSG, 10 μl samples were vigorously mixed with 0.2 μl of 2-vinylpyridine and 0.6 μl triethanolamine. After one hour, the sample was assayed as described above in the DTNB-GSSG reductase-recycling assay. According to the methods described above, standards (0.01, 0.02, 0.1, 0.2, and 1.0 mM) of total glutathione or GSSG were also assessed. The amount of total glutathione and GSSG in each sample was normalized by the protein concentration of the sample that was determined by BCA Protein Assay (Thermo Scientific, Waltham, MA, USA). The GSH concentrations were calculated from the differences between the concentrations of total glutathione and the concentrations of GSSG.

### Statistical analyses

All data are presented as mean ± SE. Data were assessed by one-way ANOVA, followed by Student-Newman-Keuls *post hoc* test. *P* values less than 0.05 were considered statistically significant.

## Results

We determined if NAD^+^ treatment may affect the oxidative stress of rotenone-treated PC12 cells. FACS-based assays using DHE as an oxidative stress probe were conducted to determine the oxidative stress of the cells. Our study has indicated that rotenone induced a significant increase in the red fluorescence intensity of ethidium, which was significantly attenuated by NAD^+^ treatment (Figure 1).

**Figure 1.**
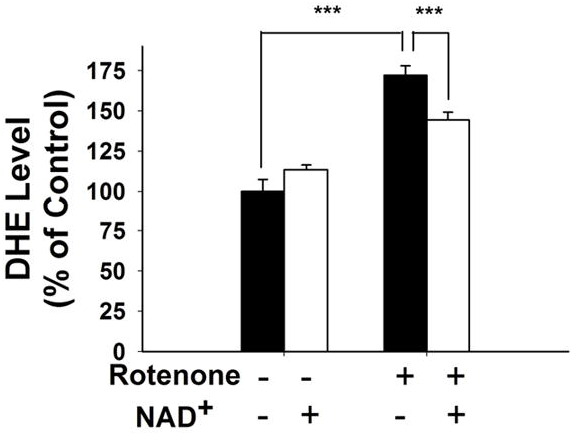
NAD^+^ treatment attenuated the rotenone-induced increase in oxidative stress in differentiated PC12 cells. Quantifications of the FACS-based determinations of the ethidium fluorescence intensity of the cells indicated that rotenone induced a significant increase in the ethidium fluorescence intensity, which was significantly attenuated by the NAD^+^ treatment. The cells were treated with 0.75 μM rotenone and 1 mM NAD^+^. The FACS-based assays were conducted 24 hours after the drug treatment. ***, *p* < 0.001.

GSH/GSSG ratio is a major index of antioxidant capacity of cells [6]. Our study determined the effects of NAD^+^ and rotenone on the GSH/GSSG ratios and the levels of GSH, GSSG and total glutathione (GSH + GSSG) in both control and rotenone-treated cells. Rotenone treatment induced significant decreases in the levels of GSH and total glutathione, which were significantly attenuated by the NAD^+^ treatment (Figures 2A and 2C). In contrast, rotenone treatment did not affect the GSSG level of the cells (Figure 2B). Rotenone treatment also induced a significant decrease in the ratio of GSH/GSSG, which was significantly attenuated by the NAD^+^ treatment (Figure 3).

**Figure 2.**
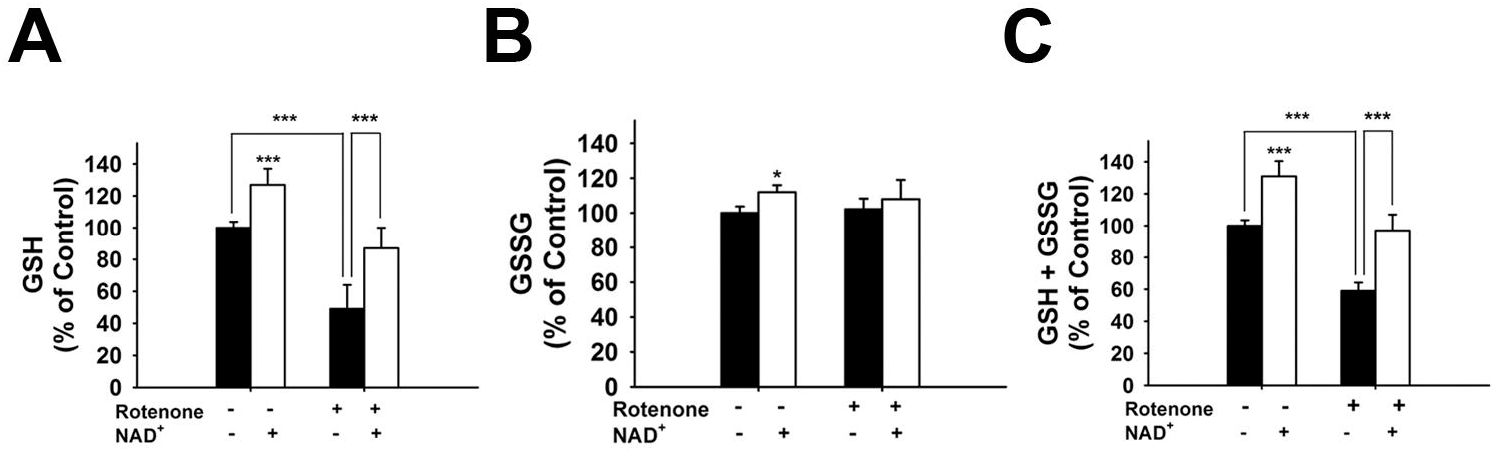
NAD^+^ treatment reversed rotenone-induced glutathione depletion in differentiated PC12 cells. (A) Rotenone induced a significant decrease in the GSH levels, which was attenuated by NAD^+^ treatment. (B) Rotenone did not affect the GSSG levels, while NAD^+^ slightly increased the GSSG levels. (C) Rotenone induced a significant decrease in the levels of total glutathione, which was blocked by NAD^+^ treatment. The cells were treated with 0.75 μM rotenone and 1 mM NAD^+^. The glutathione assays were conducted 24 hours after the drug treatment. N = 12. The data were pooled from three independent experiments. *, *p* < 0.05; ***, *p* < 0.001.

**Figure 3.**
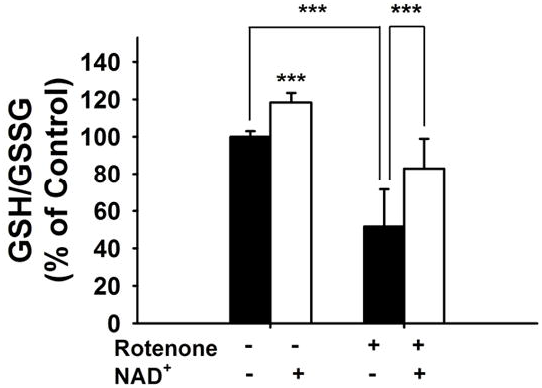
NAD^+^ treatment reversed rotenone-induced decrease in GSH/GSSG ratio in differentiated PC12 cells. Rotenone induced a significant decrease in the ratio of GSH/GSSG, which was attenuated by NAD^+^ treatment. The glutathione assays were conducted 24 hours after the drug treatment. N = 12. The data were pooled from three independent experiments. ***, *p* < 0.001.

## Discussion

The major findings of our current study include: First, NAD^+^ treatment can significantly attenuate the rotenone-induced increase in oxidative stress in differentiated PC12 cells; second, NAD^+^ treatment can attenuate the rotenone-induced decreases in the GSH and total glutathione of the rotenone-treated cells; and third, NAD^+^ treatment can attenuate the rotenone-induced decrease in the GSH/GSSG ratio of the rotenone-treated cells. Collectively, our study has provided the first evidence indicating that NAD^+^ treatment can increase directly the antioxidant capacity of cells.

NAD^+^ administration can produce profound protective effects in the animal models of aging and numerous diseases [19]. These findings, together with the observations regarding the NAD^+^ deficiency in these models, have suggested that NAD^+^ deficiency is a key common pathological factor of aging and numerous diseases [19]. In order to develop NAD^+^ administration into an effective therapeutic strategy for major diseases, it is crucial to further investigate the mechanisms underlying the beneficial effects of NAD^+^ administration. Oxidative stress play key pathological roles in aging and a number of major diseases [7, 11]. Previous studies have suggested that NAD^+^ treatment can decrease oxidative stress-induced cell death indirectly by several mechanisms, including prevention of glycolytic inhibition, mitochondrial permeability transition and nuclear translocation of AIF, as well as enhancement of SIRT1 activity [1, 2, 8]. However, it has remained unknown if NAD^+^ administration may decrease oxidative cell death by increasing directly the antioxidant capacity of the cells.

Our current study has provided the first direct evidence suggesting that NAD^+^ treatment can increase directly the antioxidant capacity of the cells exposed to oxidative stress: 1) NAD^+^ treatment can significantly increase the GSH/GSSG ratio of the rotenone-treated cells; and 2) NAD^+^ treatment can significantly attenuate the rotenone-induced increase in oxidative stress in the rotenone-treated cells. Our current finding has suggested a critical mechanism for accounting for the profound protective effects of NAD^+^ administration in multiple disease models: NAD^+^ administration can decrease oxidative stress-induced cellular damage not only by such mechanisms as prevention of glycolytic inhibition and mitochondrial alterations, but also by enhancing directly the antioxidant capacity of the cells.

There has been rapidly increasing interest on the nutritional potential of NAD^+^ [10, 14]. The finding of our current study has provided a strong evidence indicating the nutritional potential of NAD^+^: Since numerous studies have indicated age-dependent increases in oxidative stress in human body [13], nutritional supplements that can decrease oxidative stress-induced cellular damage may significantly slow the aging process. Therefore, NAD^+^ may act as a highly promising nutritional supplement, since it can enhance directly the antioxidant capacity of the cells exposed to oxidative insults.

Our latest study has indicated that NAD^+^ can also enhance directly the antioxidant capacity of the cells under basal conditions, which is mediated by SIRT2, ERK, and Nrf2 [9]. We propose that in the cells exposed to oxidative stress, NAD^+^ may also increase directly the antioxidant capacity of the cells by modulating the activities of SIRT2, ERK, and Nrf2. Future studies are warranted to test the validity of this proposal.

## Acknowledgment

The authors would like to acknowledge the financial support by a Major Special Program Grant of Shanghai Municipality (Grant # 2017SHZDZX01) (to W.Y.), and a Major Research Grant from the Scientific Committee of Shanghai Municipality #16JC1400500 and #16JC1400502 (to W.Y.).

